# DisenTE: Sparse Pattern–Context Modeling for Interpretable Translation-Efficiency Matrix Completion

**DOI:** 10.64898/2026.09.23.753721

**Authors:** Huo Miaozhe, Yu Yingying, Li Shiying

## Abstract

Partially observed object-by-context matrices arise across data-rich science, where dominant object effects can obscure smaller but informative context-dependent variation. We study this problem in a translation-efficiency atlas of 9,494 5′ UTRs across 78 cellular and tissue contexts. We present DisenTE, a sequence-conditioned neural model that combines separate sequence and context branches with a sparse low-rank pattern–context channel. Each module pairs a sequence-derived activation with context-specific deployment weights, forming a dictionary whose sequence and context components can be examined separately. Under five-fold within-panel entry masking, DisenTE achieves a UTR-centered residual Spearman correlation of 0.641±0.005, compared with 0.304±0.003 for the strongest reference model. The learned dictionary retains 11 of 20 candidate modules. CTM 6 has the largest overlap with an external TOP set and a cap-proximal pyrimidine pattern; CTMs 5 and 7 also overlap the set but have purine-containing consensuses. The evidence supports CTM 6 as a TOP sequence anchor and CTMs 5 and 7 as TOP-set-associated factors. On this dataset, DisenTE improves completion over the evaluated references and provides module-level summaries of its fitted context-dependent variation.

## 1 Introduction

Many data-rich scientific studies measure the same objects under many contexts. The result is an object-by-context response matrix: rows may represent molecules, samples, or candidate designs, while columns represent cell types, experimental conditions, or environments. Measurements are often incomplete, which makes prediction a matrix completion problem. Yet the largest source of variation is frequently an object-level effect shared across contexts. A model can therefore predict the matrix well while missing the smaller variation that distinguishes how an object behaves from one context to another.

Translation efficiency provides a clear instance of this structure. Translation efficiency (TE) measures ribosome occupancy relative to the abundance of an mRNA and is estimated transcriptome-wide by combining ribosome profiling with RNA sequencing [1]. The 5′ untranslated region (5′ UTR) is the sequence before the main protein-coding region. It contains signals that influence where and how efficiently ribosomes initiate translation, including Kozak contexts, upstream open reading frames, 5′ terminal oligopyrimidine (TOP) tracts, and RNA structures [2–6]. The atlas spans cell lines, primary cells, and tissues, which we collectively call contexts. These backgrounds differ in the machinery that reads 5′ UTR signals, so one sequence can have different TE values across contexts.

The resulting atlas is a partially observed matrix *M* (*u, c*), with a 5′ UTR *u* on each row and a context *c* on each column. The mean TE of a UTR accounts for much of the matrix, while the variation around that mean describes how its translation changes across cellular contexts. One possible source of repeated structure is a regulatory sequence pattern that occurs in many UTRs and has different fitted effects across contexts. This motivates coupling matrix completion with a pattern-level representation that shares sequence information across rows and records context variation across columns.

Existing sequence models predict translation in individual assay backgrounds or across multiple cellular contexts (Section 2). Multitask prediction already permits context-specific outputs. Our focus is to organize the fitted variation into sequence-linked modules whose deployment can be examined across contexts. This requires the row representation to remain tied to sequence while the joint effect stays readable along the context axis.

DisenTE uses separate sequence and context branches together with a sparse low-rank pattern–context channel (Fig. 1). The channel is organized as Co-Translational Modules (CTMs). Each CTM combines a sequence-derived activation, a shared effect direction, and a per-context deployment weight. The resulting dictionary links sequence-derived activations to their fitted context weights

**Figure 1:**
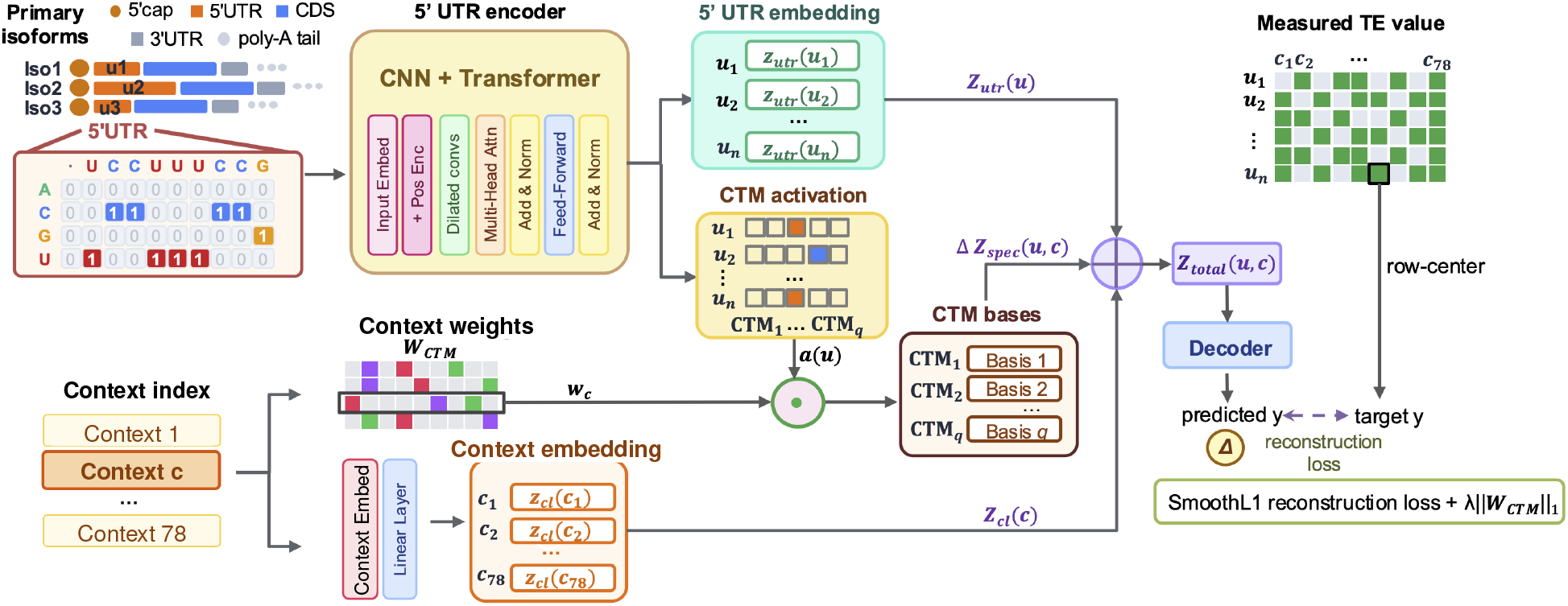
DisenTE architecture. A one-hot 5′ UTR *u* is encoded into *z*_utr_ (*u*) and CTM activations *a* (*u*), while context *c* supplies *z*_cl_ (*c*) and the row *w*_*c*_ of *W*_CTM_. The product *w*_*c*_ ⊙ *a* (*u*) is projected through CTM bases to yield Δ*z*_spec_ (*u, c*). The model decodes *z*_utr_ (*u*) + *z*_cl_ (*c*) + Δ*z*_spec_ (*u, c*) to TE and trains with row-centered smooth-*l*_1_ loss plus an *l*_1_ penalty on *W*_CTM_.

**Figure 2:**
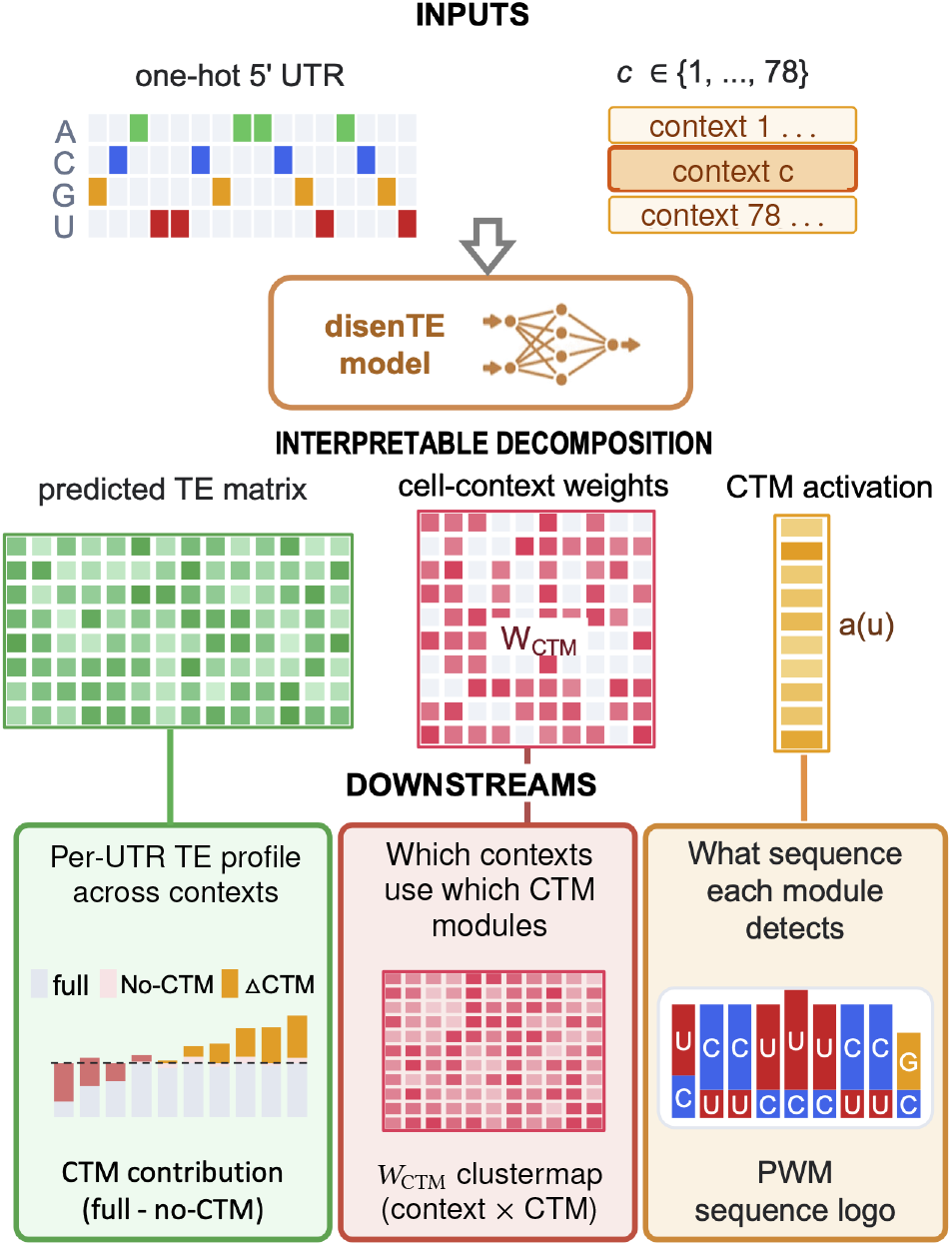
Pipeline for interpreting DisenTE. Inputs are a one-hot 5′ UTR and a context index. After fitting, the predicted TE matrix, *W*_CTM_, and CTM activations *a* (*u*) support the RPS6 full-versus-no-CTM profile, the *W*_CTM_ clustermap, and per-module PWM readout.

The main contribution is a pattern–context parameterization for sequence-conditioned matrix completion. We formulate cell-conditional TE prediction as completion of a marginal-dominated response matrix and evaluate the within-UTR variation directly. DisenTE represents that variation with a compact CTM dictionary, and the empirical study tests both sides of this claim: held-out matrix entries assess completion, while sequence attribution and an external literature-derived gene set assess possible regulatory interpretations of the fitted factors.

## 2 Related Work

### Matrix completion with side information

Low-rank factorization represents an object–context response through latent factors associated with its row and column. Factorization machines generalize this idea to pairwise interactions among observed features [10], while inductive matrix completion maps row and column side information through a low-rank bilinear operator [11]. DisenTE follows the same structural principle but derives the object features directly from sequence. Its CTM channel factorizes context-dependent variation into sequence-triggered module activations, context deployment profiles, and latent effect directions.

### Separated and compositional representations

Domain Separation Networks partition a representation into shared and domain-private components, using reconstruction and orthogonality to organize the two spaces [12]. The closest compositional precedent is CPA, which decomposes a perturbed expression profile into a basal cell-state representation and perturbation and covariate embeddings that can be recombined to predict responses [13]. CellCap later uses attention and sparse dictionary learning to decompose observed perturbation responses into transcriptional programs [14]. DisenTE applies the broad separation principle to a sequence-by-context matrix: each module couples a sequence-derived activation to a deployment profile over the measured contexts.

The available side information differs from CPA’s setting. CPA starts from measured expression states and explicit perturbation and covariate descriptors. Here, the row-side descriptor is the raw 5′ UTR sequence, whereas the columns form a fixed panel of contexts without a consistently measured molecular descriptor. DisenTE therefore uses sequence side information along the UTR axis while remaining transductive along the context axis.

### Sequence-conditioned regulation

Optimus 5-Prime predicts ribosome loading from 5′ UTRs in a reporter-assay background [7]; UTR-LM provides transferable 5′ UTR embeddings [8]. RiboNN uses full-length mRNA sequences and reading-frame information to predict cell-type-resolved TE through multitask learning. Its sequence attribution and motif analyses focus on mean TE across cell types [9]. DisenTE targets within-panel completion of UTR-centered TE variation through a sparse module parameterization linking sequence activations to context deployment weights. In adjacent regulatory genomics, Enformer predicts cell-conditional genomic tracks from DNA sequence [18].

### Reading learned sequence patterns

Regulatory sequence models are commonly interpreted with attribution methods such as Integrated Gradients [22], attribution-based motif discovery such as TF-MoDISco [23], or architectures whose filters can be summarized as motifs, including DeepBind [24] and ExplaiNN [25]. A CTM adds a context axis to this view: attribution summarizes the sequence pattern that activates a module, and the learned deployment vector summarizes where that module is used.

## 3 Method

### 3.1 Problem formulation

Throughout this section, *u* indexes a 5′ UTR, the only sequence the model sees, and *c* indexes a context. The observations form a partially filled matrix of log_2_ translation efficiencies *M* (*u, c*), with a mask marking the measured (5′ UTR, context) pairs. The task is to predict the held-out entries, a partial-matrix completion problem. In addition to raw entry prediction, we evaluate how well a model preserves variation among contexts for the same UTR.

The two views can be related through the decomposition

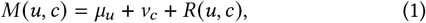

where *μ*_*u*_ is the UTR marginal, *ν*_*c*_ is the context marginal, and *R* (*u, c*) is their non-additive interaction. The UTR marginal accounts for much of the matrix variation (Section 4.8). We therefore train on the within-UTR deviation *M* (*u, c*) *μ* = *ν R* (*u, c*) by subtracting each UTR’s training mean 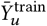 from its targets and adding that mean back at inference. This is also the quantity assessed by the UTR-centered metric in Section 4.5. The formulation preserves the context marginal while preventing the dominant row mean from deciding the score.

We model possible repeated structure across both axes with a shared pattern detector on the row side and a deployment profile on the column side. The same factors are used for completion and for summarizing fitted context-dependent patterns.

### 3.2 Sparse pattern–context factorization

DisenTE implements this idea with three paths in a joint latent predictor (Fig. 1). It predicts log_2_ TE by applying a shared decoder to their sum,

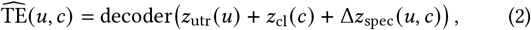

where *z*_utr_ *u* represents the 5′ UTR alone, *z*_cl_ *c* represents the context alone, and Δ*z*_spec_ *u, c* is computed jointly from sequence and context. The disjoint inputs make the first two paths recognizable as row- and column-side components. The shared nonlinear decoder can still combine all three terms, so Equation (2) is a structured latent parameterization rather than an additive decomposition of the final prediction.

The joint path uses a low-rank bilinear form,

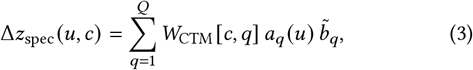

where *a*_*q*_ *u* 0 is a sequence-derived activation, 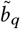 a unit-norm effect direction, and *W*_CTM_ *c, q* a per-context weight; in matrix form 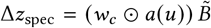. Each rank-one term is active when a sequence-derived scalar *a*_*q*_ (*u*) is paired with the context weight *W*_CTM_ [*c, q*]; its signed magnitude is projected along 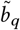. The low-rank form supplies an explicit, removable route for pattern–context modulation and connects directly to the matrix factorization view in Section 2.

We call each of the *Q* terms in Equation (3) a Co-Translational Module (CTM). A convolutional detector *a*_*q*_ provides the UTR-side activation, a latent direction 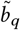 represents its effect in the shared prediction space, and a column *W*_CTM_ [·, *q*]records deployment across contexts. An *l*_1_ penalty on*W*_CTM_ encourages a compact set of active modules. Normalizing each 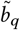 to unit norm removes the per-module scale ambiguity between the deployment weights and the effect directions; sign and permutation symmetries remain. Active columns and their context profiles can be read directly from *W*_CTM_; the sequence evidence associated with each module is summarized separately from its activation map (Section 4.9).

### 3.3 Implementation

A shared encoder produces the sequence representation and module activations. Each 5′ UTR is tokenized and right-anchored (Section 4.2), then passed through three dilated convolutional layers, which provide a local receptive field for short cap-proximal motifs, followed by two Transformer layers [15], implemented here with pre-layer normalization, that integrate information across the 5′ UTR.

The encoder emits *z*_utr_ (*u*) by concatenating mean- and max-pooled hidden states over unmasked positions. A parallel convolutional head uses masked max pooling to produce the CTM activations *a* (*u*). Its position-wise activation map is retained for the later extraction of per-CTM position-weight matrices.

The context branch supplies *z*_cl_ (*c*) from a context embedding and a linear projection. It also supplies the row *w*_*c*_ = *W*_CTM_ [*c*, :]of the cell-context weight matrix. This matrix is initialized to zero, leaving the CTM channel inactive until the reconstruction loss recruits it. A two-layer perceptron decodes the summed representation into a scalar TE prediction. The hidden width is 128, and the model starts with *Q* = 20 candidate CTM channels. Section 4.4 gives the remaining dimensions and optimization settings.

## 4 Experiments

The evaluation asks two questions. Can the model complete held-out entries after the dominant UTR marginal has been removed? Does the low-rank channel organize the remaining variation into modules that are supported by both sequence and gene-set evidence? Component ablations relate these questions by measuring how much each model path contributes to the completion score.

### 4.1 Dataset and preprocessing

We use the released log_2_ TE values from the human compendium of the RiboNN ribosome-profiling resource [9] as our regression target. Each entry averages measurements for the same transcript within a context. The 78 contexts include cell lines, primary cells, and tissues. For each gene we keep one 5′ UTR, drawn from the MANE Select transcript where available and from the APPRIS principal isoform otherwise. A 5′ UTR is retained only if its length lies in [30, 512] nt and the corresponding gene is covered by at least 30 contexts in the RiboNN TE matrix. Near-duplicate 5′ UTRs are collapsed by 5-mer Jaccard ≥0.7 within ±10% length bins. The resulting atlas comprises 9,494 5′ UTRs across 78 cellular and tissue contexts, with 661,457 observed (5′ UTR, context) TE pairs (89.3% coverage of the 5′ UTR–context matrix); the median 5′ UTR is 147 nt long. Each observed pair is represented by its 5′ UTR sequence and context index.

### 4.2 Sequence encoding

Each 5′ UTR is tokenized over an alphabet of six symbols (the four nucleotides, an unknown symbol, and a padding symbol), with thymine mapped to uracil and any unrecognized character mapped to the unknown symbol. The resulting token sequence is placed flush against the right edge of a window of length *L* = 512 by left-padding, so that the position immediately upstream of the main start codon occupies a common coordinate across all 5′ UTRs. A binary attention mask of shape (*B, L*) marks real and padded positions and excludes the latter from all subsequent computations.

### 4.3 Cross-validation splitting protocols

We evaluate DisenTE under two splitting regimes, both operating at the 5′ UTR level, with the same fold memberships shared across every model and ablation.

The headline benchmark uses a within-panel five-fold entry-masking protocol. For each 5′ UTR, its observed contexts are partitioned round-robin into five buckets under a fixed seed; in fold *k*, the *k*-th bucket forms the test set, the next bucket forms the validation set, and the remaining three buckets form the training set. All 78 context identities remain represented in every training fold through other 5′ UTRs. Across the five folds, the pooled test set contains 661,457 entries.

For the interpretation analyses we train a single reference model under the all-data regime, in which the 661,457 observed pairs are randomly partitioned into 95% training and 5% validation. The validation portion supports training diagnostics and checkpoint selection; module analyses use the fitted reference model.

### 4.4 Training objective and optimization

The per-5′ UTR mean-centering of the target (Section 3.1) is applied under both the five-fold benchmark and the all-data regime, so the model is fit to the cell-conditional deviation throughout. The training objective combines a smooth-*l*_1_ Huber reconstruction loss between the predicted and observed TE with an *l*_1_ penalty on the cell-context weight matrix,

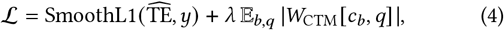

in which the expectation is taken over the batch. With *λ* = 0.10, the fitted all-data model retains 11 active CTM channels on the all-data run. Parameters are optimized with AdamW (learning rate 7 × 10^−4^, weight decay 10^−4^), batch size 128, a cosine-annealing schedule over the full training horizon, and gradient-norm clipping at 1.0. We train the benchmark and ablation models for 20 epochs and the all-data model for 25 epochs; the checkpoint with the highest validation per-context ρ_res_ is retained. All experiments use PyTorch 2.11 on Python 3.10, with training and inference on a single NVIDIA RTX 4090 GPU.

### 4.5 Evaluation metrics

For a test partition we let *P* and *Y* denote the per-pair predicted and observed TE values and write 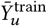 for the per-5′ UTR mean computed on the training pairs alone. The residual at a pair is its deviation from this per-5′ UTR baseline.

We report the residualized Spearman correlation ρ_res_, computed between predictions and observations after each is centered on the per-5′ UTR mean, together with the raw Spearman ρ. Because any model can obtain a high raw correlation from the dominant UTR marginal, ρ_res_ is our primary metric. It discounts this baseline and measures the ranking of context-dependent deviations. For cross-model comparison we use a symmetric residual, in which the predictions and observations are each recentred on their own per-5′ UTR means computed on the test set, 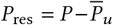 and 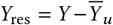. This eliminates a calibration artefact that would otherwise penalize models recovering the per-5′ UTR mean to different precision.

Our primary per-context criterion is the cell-conditional Spearman: for each context *c* with at least ten test pairs we compute ρ_res,*c*_ = Spearman (*P*_res,,*c*_, *Y*_res,,*c*_) and average across contexts. This is also the metric on which validation checkpoints are selected.

For each fold, we compute the metrics after pooling all held-out pairs in that fold. The five disjoint test partitions contain *n* = 661,457 pairs in total, and table entries report the mean and standard deviation of the five fold-level statistics. We also report ρ_res_ on nested strata of 5′ UTRs ranked by cross-context TE standard deviation (the top 5, 10, 20, and 50%, and all 5′ UTRs). For the ablation figure we additionally compute a per-5′ UTR Spearman across held-out contexts and report the median within each stratum.

### 4.6 Baselines

We compare DisenTE against five baselines that share its splitting protocols and target conventions, so that the rows of the stratified comparison panel (Table 1) can be read directly. Each sequence reference is paired with context identity. The Ridge models concatenate sequence features with a 78-dimensional context one-hot vector; the neural models add a shared context tower consisting of an 8-dimensional embedding and a linear projection to a scalar. Optimus 5-Prime is trained under this paired-input setting, whereas UTR-LM is used as a frozen sequence encoder as detailed below. The two neural baselines use AdamW and a cosine-annealing learning-rate schedule.

**Table 1:** Fold-level pooled UTR-centered residual Spearman *ρ*_res_ on the five-fold entry-masking benchmark, stratified by 5′ UTR cross-context TE standard deviation. Values are mean ± s.d. over folds.

| Model | Top 5% | Top 10% | Top 20% | Top 50% | All 5' UTRs |
| --- | --- | --- | --- | --- | --- |
| Two-way mean baseline<sup>a</sup> | 0.196 ± 0.007 | 0.212 ± 0.006 | 0.243 ± 0.006 | 0.274 ± 0.006 | 0.300 ± 0.003 |
| 6-mer sequence fusion model | 0.195 ± 0.008 | 0.212 ± 0.006 | 0.245 ± 0.006 | 0.277 ± 0.006 | 0.304 ± 0.003 |
| Translation-rule fusion model<sup>b</sup> | 0.195 ± 0.008 | 0.212 ± 0.006 | 0.244 ± 0.007 | 0.277 ± 0.006 | 0.304 ± 0.003 |
| Optimus 5-Prime [7] fusion model | 0.197 ± 0.009 | 0.213 ± 0.006 | 0.245 ± 0.006 | 0.277 ± 0.006 | 0.304 ± 0.003 |
| UTR-LM [8] fusion model | 0.211 ± 0.011 | 0.219 ± 0.006 | 0.244 ± 0.007 | 0.265 ± 0.007 | 0.281 ± 0.003 |
| DisenTE (ours) | **0.687 ± 0.008** | **0.672 ± 0.002** | **0.669 ± 0.004** | **0.663 ± 0.005** | **0.641 ± 0.005** |

<sup>a</sup> Predicts $\bar{Y}_u^{\text{train}} + \bar{Y}_c^{\text{train}} - \bar{Y}^{\text{train}}$ .
<sup>b</sup> Ridge on the 13 hand-crafted 5' UTR features of Appendix A.
Each sequence representation is paired with context identity as described in Section 4.6.

- **Two-way mean baseline**. A non-learned reference that predicts 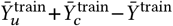 for every (5′ UTR, context) pair,
- the additive combination of the per-5′ UTR and per-context training marginal means. This is the strongest marginal reference among the global, per-5′ UTR, per-context, and context-mean variants.
- **6-mer sequence fusion model**. A per-fold Ridge regression (*α* = 1.0) on the concatenation of the normalized 6-mer count vector of the 5′ UTR and the context one-hot vector, representing the contribution of low-order *k*-mer composition.
- **Translation-rule fusion model**. A per-fold Ridge regression on the 13 hand-crafted 5′ UTR annotations of Appendix A (Kozak strength, upstream-AUG counts, closed-uORF descriptors, an IRES-hit indicator, etc.) concatenated with a context one-hot vector, representing curated translation-initiation features.
- **Optimus 5-Prime fusion model** [7]. We adapt the convo-
- lutional backbone of Sample et al. Our version uses three Conv1d layers with kernel size 8 and 120 channels, each followed by ReLU and dropout, flattened to a two-layer fully connected head, operating on the rightmost 256 nucleotides of the 5′ UTR, fused with the context tower as above.
- **UTR-LM** [8]. In this fusion model, the right-anchored 256-nucleotide 5′ UTR is passed through the frozen six-layer UTR-LM Transformer of Chu et al.; the 128-dimensional class-token embedding is fed to a small head of one linear layer, a ReLU, dropout, and a final linear projection, fused with the context tower as above. The UTR-LM weights are not updated during training; only the head and the context tower are learned.

In addition to the five baselines above, we evaluate four architectural ablations of DisenTE, each removing one component at a time. Removing the *cell-context module dictionary* sets Δ*z*_spec_ ≡ 0 and measures the contribution of the sparse bilinear pattern × context channel. Removing the *additive context embedding* sets *z*_cl_ ≡ 0 and measures the contribution of the context-only path. Removing the *Transformer encoder* retains only the dilated CNN front and tests the contribution of global integration to motif detection. Removing the *dilated convolutional encoder* retains only Transformer layers and tests the contribution of local receptive fields. All ablations are trained on the same fold splits and hyperparameters as the full model; ablation results and their interpretation are reported in Section 4.11.

### 4.7 Module analysis protocol

The interpretation analyses all operate on the all-data reference model. For each CTM we take its top-50 most strongly activating 5′ UTRs, the 50 with the highest per-CTM activation *a*_*q*_ (*u*), as the module’s representative set. Integrated gradients [22] trace its max-pooled activation back to each 5′ UTR position using 50 integration steps. Aggregating these scores over the top-50 gives the position-weight matrix and information content reported in Appendix B.

For the anchor module CTM 6, we compare its top-50 with the remaining 5′ UTRs along three axes (Fig. 4): UTR length, a binary cap-proximal C + pyrimidine 5′ TOP regex, and m^6^A RRACH density per kb. The continuous variables use a Mann–Whitney test with rank-biserial *r*; the binary regex uses a Fisher exact test. External support comes from a 79-gene literature TOP set containing ribosomal proteins and canonical translation factors. Using the full set of 9,494 retained UTRs as the comparison universe, we test each CTM’s top-50 overlap with Fisher’s exact test and a 10,000-draw null of random size-50 sets (expected overlap 0.42 hits), with Bonferroni correction over the 11 surviving CTMs. Gene-family annotations summarize the top-50 sets in Fig. 3 but are not model inputs.

**Figure 3:**
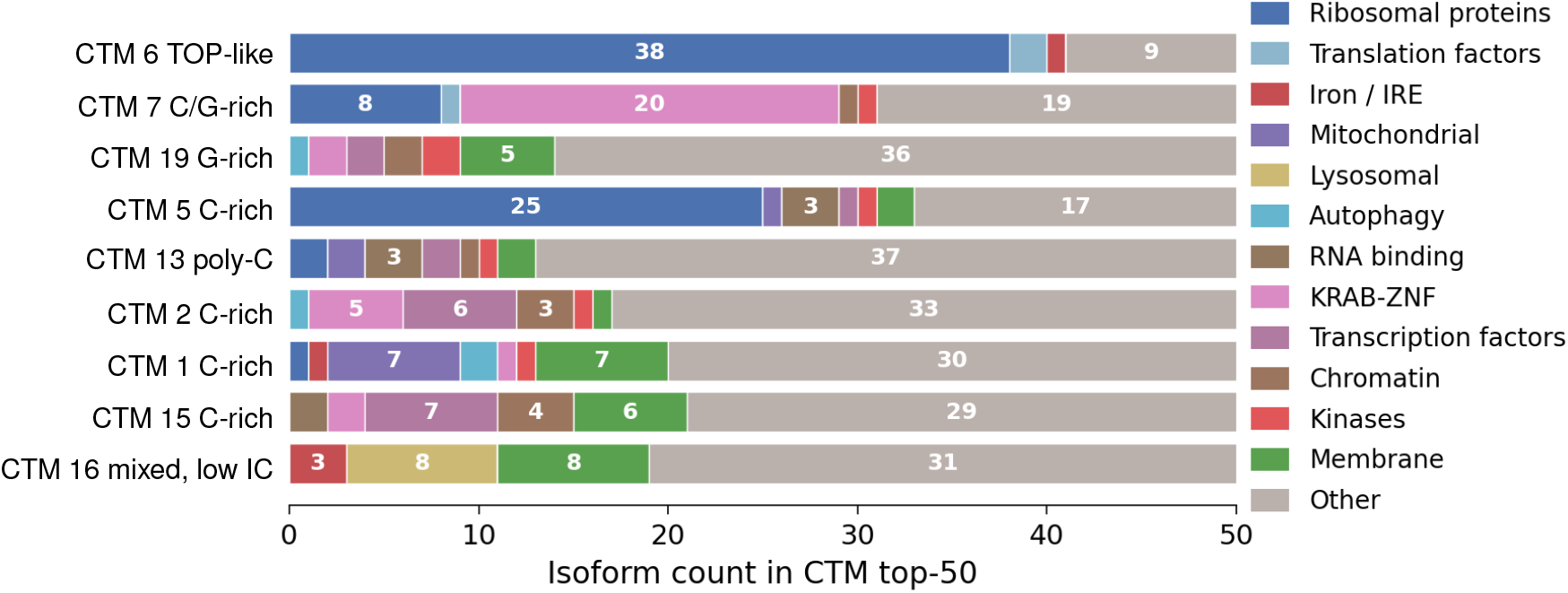
Gene-family composition of CTM top-50 5′ UTRs. Row labels describe PWM sequence composition, matching Table 4; colors report gene functional classes. TOP-set support is quantified for CTMs 5–7, with CTM 6 dominated by ribosomal-protein 5′ UTRs.

**Figure 4:**
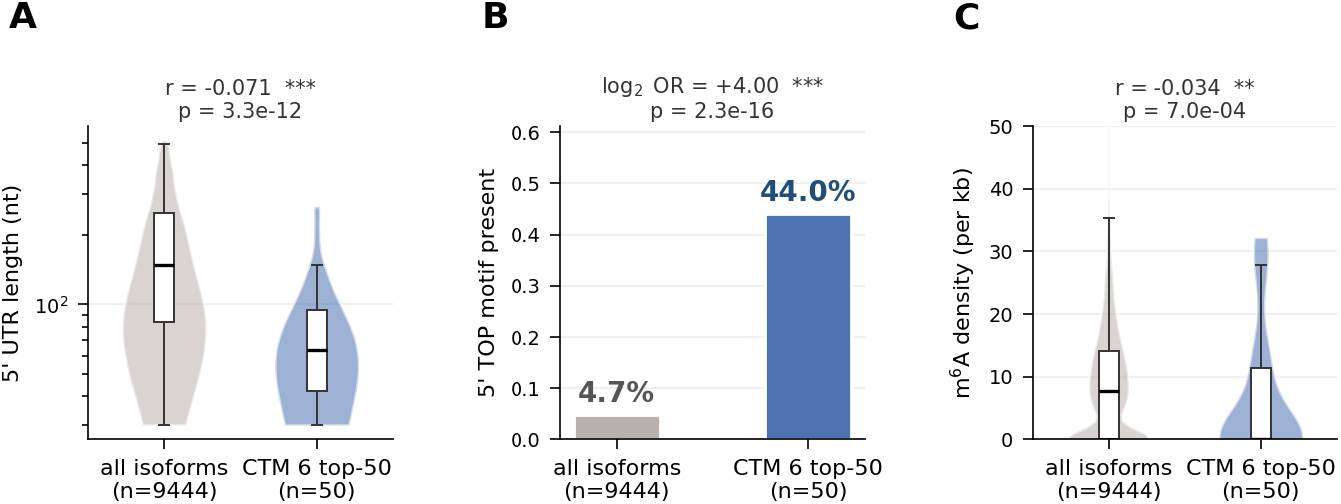
CTM 6 anchor evidence. The CTM 6 top-50 5′ UTRs (blue) are compared with the other 9,444 5′ UTRs (grey) by (A) 5′ UTR length, (B) cap-proximal 5′ TOP regex, and (C) RRACH density. Effect sizes and *p* values are shown above panels.

For the RPS6 case study, we decompose each per-context prediction as 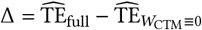, isolating the contribution of the bilinear CTM channel by zeroing the cell-context weights.

### 4.8 Within-panel matrix completion

On the five-fold entry-masking split (Section 4.3), the additive marginals account for most of the raw signal but little of the within-UTR variation. The two-way mean baseline reaches a raw Spearman of 0.772 but ρ_res_ = 0.300 after UTR centering. The four sequence-fusion references fall in a narrow range from 0.281 to 0.304 on the same criterion, including the pretrained UTR-LM representation and the adapted Optimus 5-Prime CNN (Table 1). DisenTE reaches ρ_res_ = 0.641±0.005 on all UTRs.

The variability strata show where the difference is largest. Among the top-5% most variable UTRs, DisenTE reaches ρ_res_ = 0.687, whereas the reference models reach at most ρ_res_ = 0.211. DisenTE’s score rises as the analysis narrows to more variable UTRs; every reference score falls. The gain is therefore concentrated in variation across contexts for the same UTR rather than in the row marginal alone. Section 4.11 separately measures the contribution of the bilinear CTM path and the additive context path.

### 4.9 Evidence across the learned modules

The all-data reference model retains 11 CTMs (Section 4.11), nine of which are summarized by gene-family composition in Fig. 3. CTMs 5–7 are the only modules with significant overlap with the external TOP set, and CTM 6 is the most concentrated member. We therefore base the biological interpretation below on these measured overlaps and the corresponding sequence summaries. The TOP regulon contains ribosomal proteins and other components of the translation apparatus [4, 5, 19].

The external-set results identify CTMs 5–7 as three factors whose top activations are enriched for known TOP transcripts (Table 2). Their top-50 sets contain 26, 44, and 11 TOP genes, respectively, with Fisher exact *p <* 10^−42^, *p <* 10^−90^, and *p <* 10^−12^. A 10,000-draw permutation null has an expected overlap of 0.42, a maximum of 5, and a 99.9th percentile of 3. The sequence evidence distinguishes the three modules. Classical TOP motifs begin with a cap-adjacent C followed by an uninterrupted tract of 4–14 pyrimidines [5]. CTM 6 has the fully pyrimidine-rich consensus CUCUCCC and the highest information content, 2.71 bits, consistent with this definition and with its stronger gene-set concentration. CTM 5 (CGCUCCC, 1.48 bits) and CTM 7 (CGGUCCC, 1.12 bits) contain purines; their TOP evidence therefore comes from transcript-set enrichment rather than a classical TOP consensus. CTM 5’s C-rich composition is compatible with the C-rich binding preference of PCBP1 [20], which has demonstrated translation-regulatory roles [21]. This motivates PCBP1 as a candidate reader, but neither the short PWM nor these background studies establish PCBP1 binding to CTM 5 targets.

**Table 2:** Overlap of CTM top-50 5′ UTRs with a 79-gene literature 5′ TOP set.

| CTM | Class I<sup>a</sup> | Class II<sup>a</sup> | Total | Fisher exact $p$ |
| --- | --- | --- | --- | --- |
| 5 | 26 | 0 | 26/50 | $< 10^{-42}$ |
| 6 | 41 | 3 | 44/50 | $< 10^{-90}$ |
| 7 | 9 | 2 | 11/50 | $< 10^{-12}$ |

<sup>a</sup> Class I = RPL / RPS ribosomal proteins; Class II = canonical translation factors. Literature TOP set assembled from [4, 5, 19].
Random expectation = 0.42 hits per top-50; a 10,000-draw permutation null caps the random hit count at 5. All three reported overlaps remain significant after Bonferroni correction over the 11 surviving CTMs.
Eight other PWM-anchored CTMs are omitted: six with zero hits (CTM 2, 8, 10, 15, 16, 19) and two non-significant (CTM 1: 1/50, $p = 0.34$ ; CTM 13: 2/50, $p = 0.06$ ).

We examine CTM 6 in greater detail because it has the largest TOP-set overlap and the highest information content among the displayed modules. Its top-50 UTRs have shorter 5′ UTRs (median 63.5 vs 147.5 nt), a cap-proximal C + pyrimidine regex hit at 44.0% versus 4.7% in the rest of the atlas (Fisher exact log_2_ OR = +4.00), and lower m^6^A RRACH density (median 0 vs 7.8 per kb; Fig. 4). The 44 /50 literature-set overlap is 105.8 ×the random size-50 expectation. Together with the integrated-gradients consensus, these sequence, transcript, and gene-set views give CTM 6 the strongest TOP-related support among the fitted modules.

The three displayed CTM 6 top-50 UTRs span a Class II translation factor (EEF1A1), a Class I small-subunit protein (RPS2), and a Class I large-subunit protein (RPL13A; Fig. 5). Each example has cap-proximal attribution, consistent with the regex summary in Fig. 4B and the consensus PWM in Appendix B.

**Figure 5:**
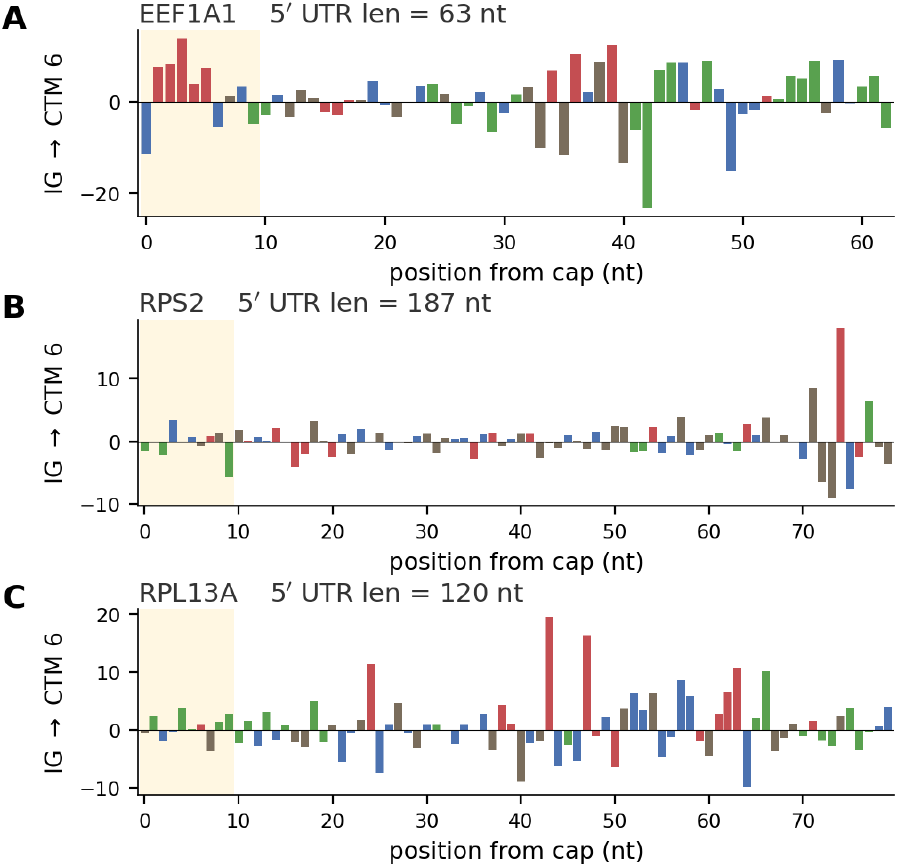
CTM 6 integrated-gradient maps for three TOP-class 5′ UTRs: (A) EEF1A1, (B) RPS2, and (C) RPL13A. Bars are per-position IG scores colored by nucleotide; the yellow band marks cap-proximal positions 0–10.

### 4.10 Per-UTR decomposition case study

In the all-data interpretation fit, DisenTE reproduces the measured context TE profile of the canonical 5′ TOP example RPS6 closely (Pearson *r* = 0.97, Spearman ρ = 0.98 across all 78 contexts; Fig. 6A). This case study describes the fitted CTM decomposition; the five-fold benchmark in Section 4.8 provides the held-out performance estimate. The full-versus-no-CTM difference in Fig. 6B varies across contexts and changes sign, rather than acting as a constant UTR-level offset. The positive side includes K562, leukemia, breast-cancer, and melanoma lines, while the negative side includes muscle, primary-neuron, and BJ-fibroblast contexts.

**Figure 6:**
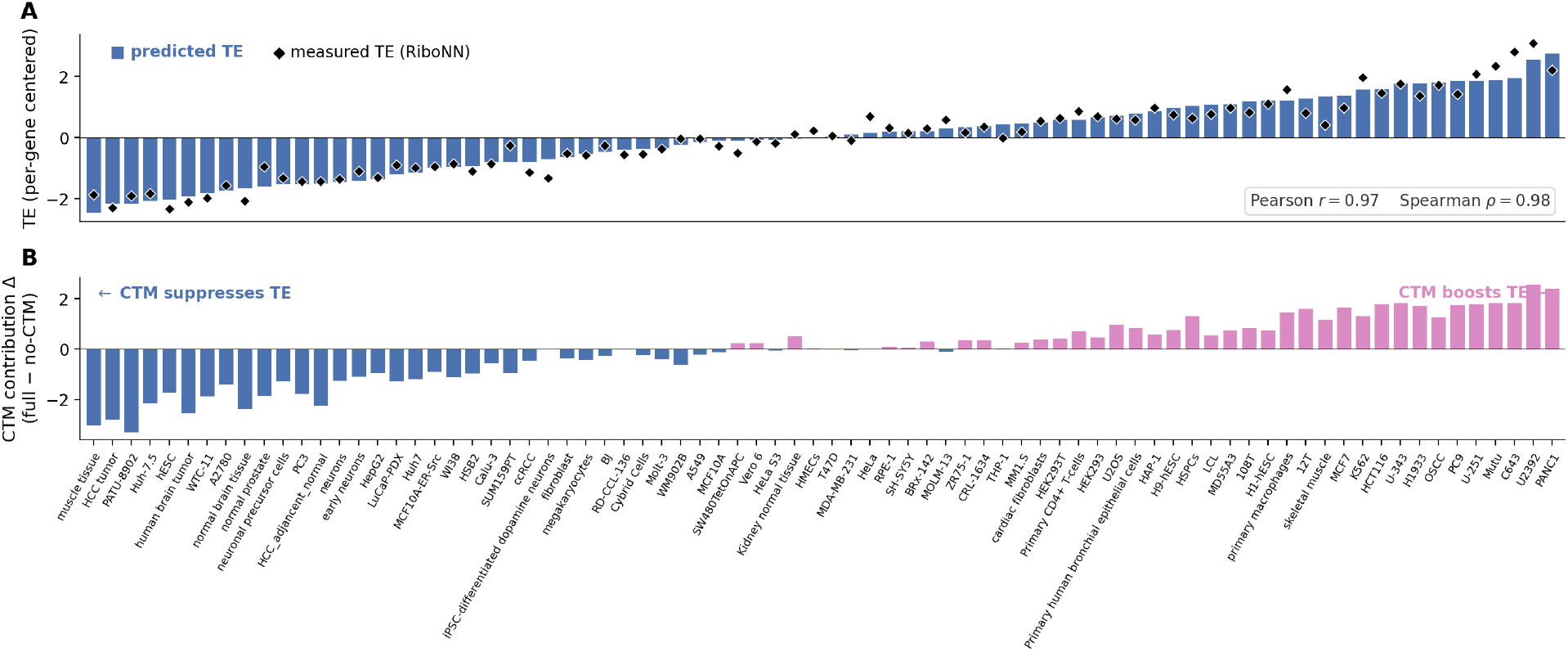
RPS6 per-context decomposition from the all-data reference fit. (A) Predicted TE (blue bars) and measured TE (black diamonds) across 78 contexts. (B) CTM contribution 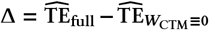. Contexts are sorted by panel A and reused in panel B.

### 4.11 Contribution of model components

Each ablation retrains DisenTE with one component removed under identical splits and hyperparameters; Figure 7 reports the change in pooled UTR-centered residual Spearman.

**Figure 7:**
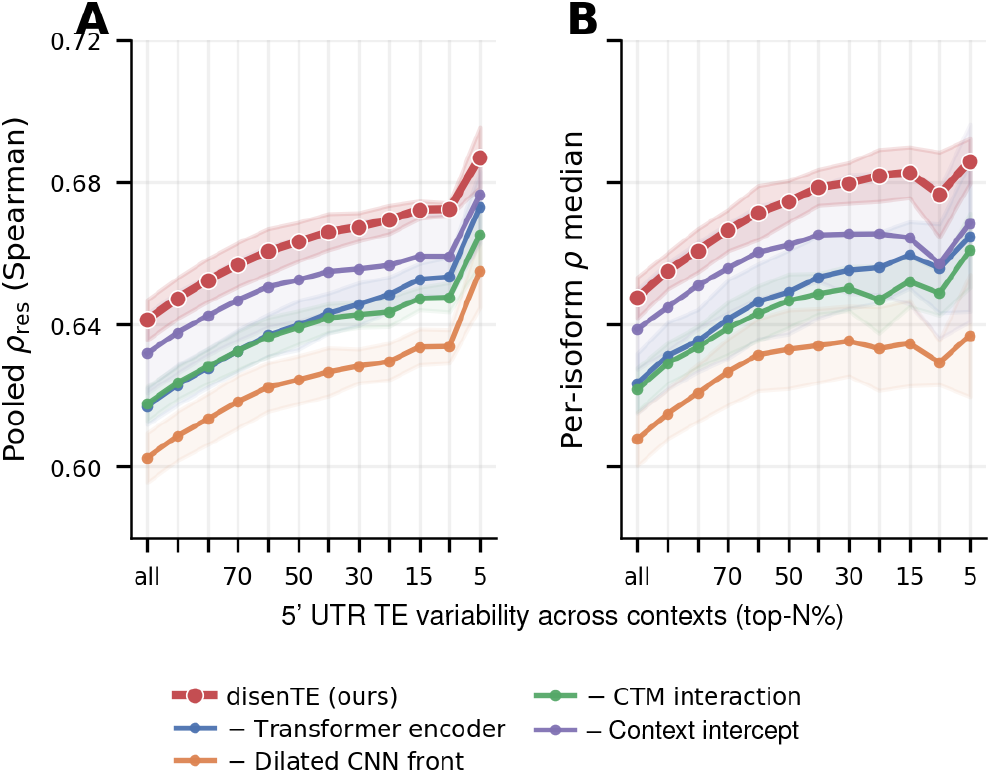
Architectural ablation on the five-fold benchmark across 5′ UTR variability strata. Curves show five-fold means with shaded 1±s.d. (A) Pooled *ρ*_res_; (B) median per-5′ UTR Spearman

The two encoder components, the dilated CNN front for local pattern detection and the Transformer for global integration, both contribute to encoding the 5′ UTR. Removing them reduces pooled ρ_res_ by 0.038 and 0.024, respectively. The dilated CNN front produces the larger ablation drop on this benchmark; removing the Transformer produces the smaller of the two encoder-component changes.

Removing the joint CTM path Δ*z*_spec_ reduces pooled ρ_res_ by 0.024, compared with 0.009 after removing the context-only path *z*_cl_ (Fig. 7). On this benchmark, removing the joint pattern × context path therefore produces the larger of the two observed drops.

The fitted dictionary contains 11 surviving channels from the 20 candidates (defined as max_*c*_ |*W*_CTM_ [*c, q*]| *>* 0.05; the remaining nine columns fall below this threshold, eight of them strictly below 10^−3^). Thus, 45% of the candidate channels are inactive under the stated threshold. The clustermap of the surviving *W*_CTM_ block (Fig. 8) groups the 78 contexts by their fitted CTM weights. Each surviving channel has its own deployment profile, and the nine displayed modules have different gene-family compositions (Fig. 3). We use *dictionary* to denote this fitted collection of module profiles.

**Figure 8:**
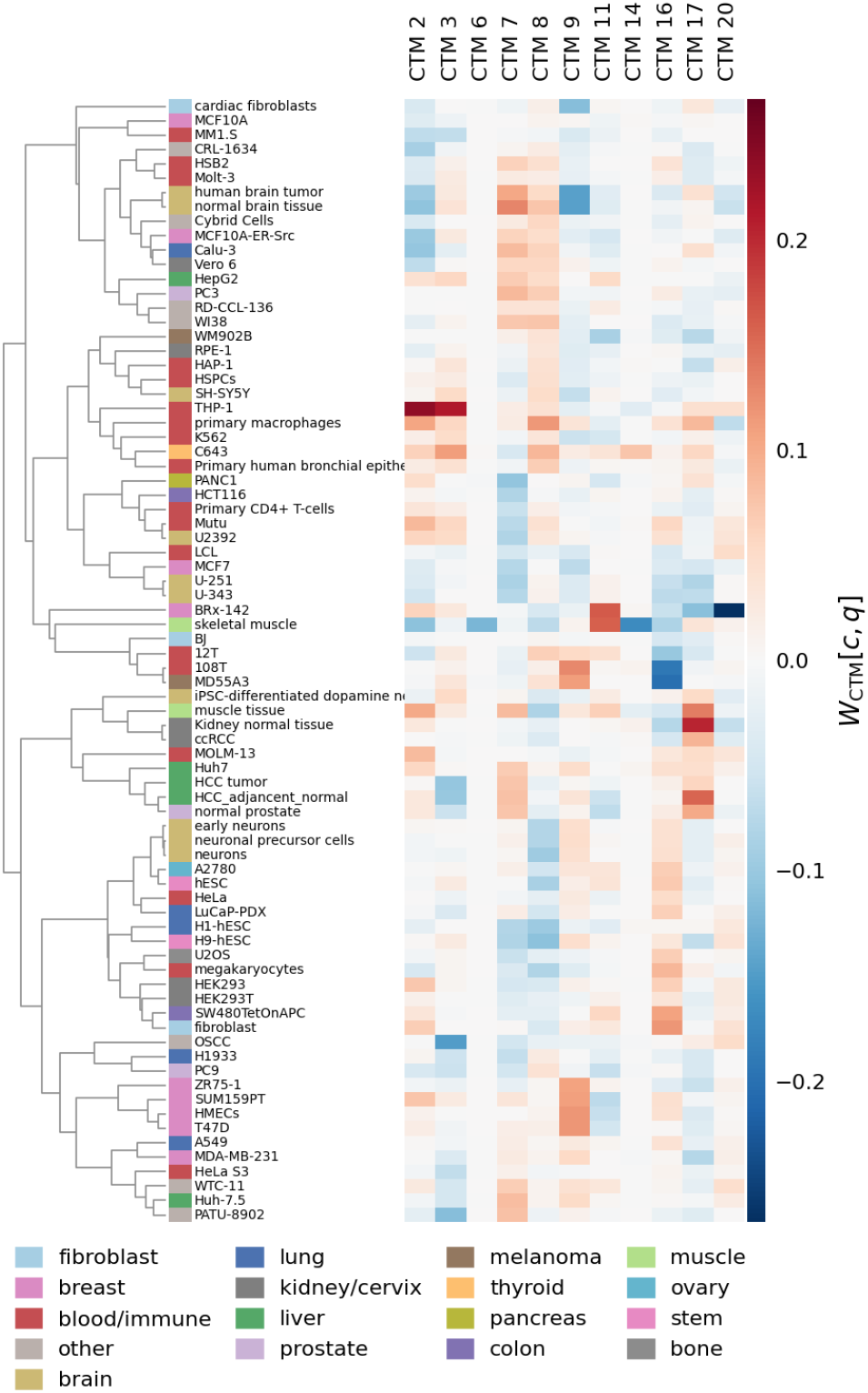
*W*_CTM_ clustermap for the all-data reference model, restricted to the 11 surviving CTM columns. Rows are the 78 contexts clustered by their surviving CTM weights; the left strip gives context group annotations for visualization.

## 5 Discussion

The gap between the raw and UTR-centered scores is central to this task. A two-way marginal model reaches a raw Spearman correlation of 0.772, showing that much of the atlas can be predicted from UTR and context averages. Its residual score is only 0.300, however, because those averages say little about which UTRs change in which contexts. DisenTE reaches 0.641±0.005 on the centered criterion, and the advantage grows among UTRs with greater cross-context variation. The main gain therefore lies in the smaller conditional structure that the raw matrix score obscures.

The architecture gives that structure a dedicated route. Separate UTR and context paths retain the two marginals, while the bilinear CTM path couples sequence-triggered activations to context weights. Removing this path lowers the pooled score by 0.024, compared with 0.009 after removing the context-only path. On this ablation benchmark, the joint pattern–context path therefore has the larger measured effect. The shared decoder remains nonlinear, so the three paths are a structured latent parameterization rather than an exact additive partition of TE.

CTM 6 is supported at both the transcript-set and sequence levels: its top activations are concentrated in known TOP transcripts, and its cap-proximal pyrimidine consensus matches the classical TOP definition. CTMs 5 and 7 also enrich for known TOP transcripts, but their consensuses contain purines. We therefore treat them as TOP-set-associated modules rather than additional TOP motif recoveries. This distinction keeps transcript composition separate from sequence identity while retaining each module’s fitted context deployment profile.

From a matrix-completion perspective, the model lies between standard transductive factorization and fully inductive completion. Raw sequence provides side information for every UTR, including a row whose entries are partly hidden, whereas the context representation is learned from its observed entries. This asymmetry matches the available atlas: the UTRs have a common descriptor, but comparable molecular profiles are not available for every column. It also explains why the current benchmark masks entries rather than entire contexts.

The context axis is transductive: every test identity appears elsewhere in training and is represented by an index embedding. The model therefore addresses unmeasured entries among the 78 observed contexts. Extending the context axis would require comparable molecular descriptors across contexts.

The learned modules are best read as structured hypotheses about the fitted response matrix. Their sequence component is obtained by attribution, and their biological label is strongest when an external gene set supplies an additional anchor. The right-anchored encoding also places the cap-proximal boundary next to padding. Encoder designs that model the cap boundary explicitly could sharpen future sequence-level analyses.

## 6 Conclusion

DisenTE casts cell-conditional translation efficiency as completion of a marginal-dominated UTR-by-context matrix. Its sparse low-rank CTM path links sequence-triggered activations to context deployment profiles inside a joint predictor. On the five-fold entry-masking benchmark, the model reaches a UTR-centered residual Spearman of ρ_res_ = 0.641±0.005, compared with 0.304±0.003 for the strongest reference. The learned dictionary retains 11 of 20 candidate modules. CTMs 5–7 have the strongest overlaps with the external TOP set, and CTM 6 supplies the clearest TOP-related sequence anchor. Together, the completion and module analyses indicate that an explicit pattern–context model can predict within-object variation while exposing module-level sequence and context summaries of the fitted context panel.

## 7 Code and Data Availability

The model code, configurations, and trained checkpoints are available at anonymous.4open.science/r/disente-code-B22E. The bench-mark uses the released RiboNN TE matrix [9] and GENCODE v38 5′ UTR sequences [16].

### A Translation-rule fusion model features

The Translation-rule fusion model of Section 4.6 uses the 13 features summarized in Table 3. Each feature is standardized within the training fold before fitting the Ridge model.

**Table 3:**
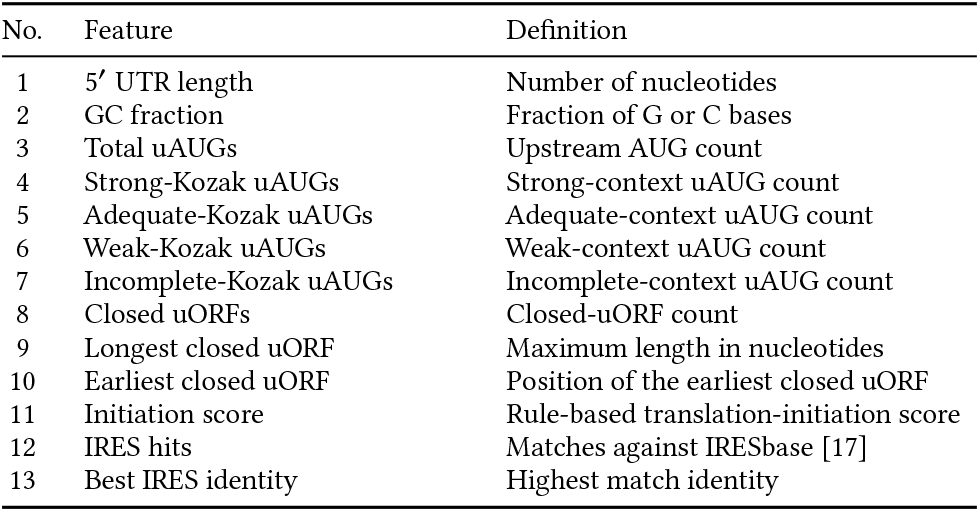
Features used by the Translation-rule fusion model.

| No. | Feature | Definition |
| --- | --- | --- |
| 1 | 5' UTR length | Number of nucleotides |
| 2 | GC fraction | Fraction of G or C bases |
| 3 | Total uAUGs | Upstream AUG count |
| 4 | Strong-Kozak uAUGs | Strong-context uAUG count |
| 5 | Adequate-Kozak uAUGs | Adequate-context uAUG count |
| 6 | Weak-Kozak uAUGs | Weak-context uAUG count |
| 7 | Incomplete-Kozak uAUGs | Incomplete-context uAUG count |
| 8 | Closed uORFs | Closed-uORF count |
| 9 | Longest closed uORF | Maximum length in nucleotides |
| 10 | Earliest closed uORF | Position of the earliest closed uORF |
| 11 | Initiation score | Rule-based translation-initiation score |
| 12 | IRES hits | Matches against IRESbase [17] |
| 13 | Best IRES identity | Highest match identity |

**Table 4:**
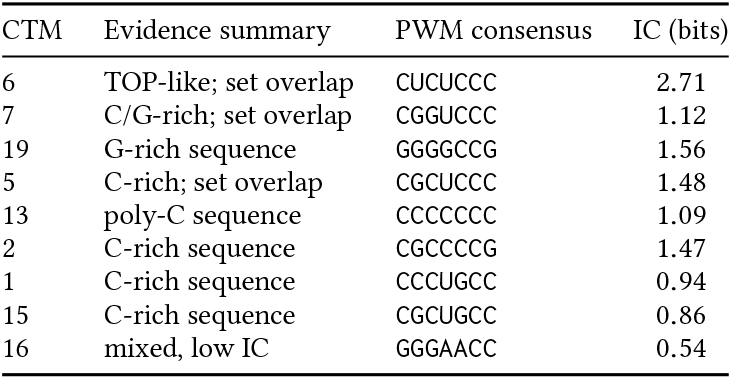
Per-CTM PWM consensus motif and total information content (in bits) for the nine surviving CTMs shown in Fig. 3.

### B Per-CTM PWM consensus and information content

Fig. 3 omits the integrated-gradients PWM consensus motif and total information content of each CTM to keep the visual focus on gene-family composition. Table 4 lists these values using sequence-composition descriptions; a seven-base PWM alone is not used to assign an RNA structure or a specific reader.

### C How to read a CTM

The CTM quantities have distinct roles. The activation *a*_*q*_ (*u*) ranks UTRs for enrichment; *W*_CTM_, [·*q*] records deployment across observed contexts; and 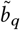 gives a latent direction, not a scalar TE effect under the nonlinear decoder. The integrated-gradient PWM summarizes sequence evidence, whereas the RPS6 ablation summarizes the complete CTM channel. Activations may overlap across UTRs; sparsity acts on deployment weights, and top-50 labels summarize activation tails.

### Ethical Considerations

The study uses public, aggregate measurements from cell lines, primary cells, and tissues, together with public transcript annotations; it contains no individual-level personal data. The predicted entries and CTM labels are intended for research prioritization followed by experimental validation. Releasing the preprocessing and evaluation code supports reproducible assessment of these uses.

